# Use of Avian Adeno-Associated Virus for the Delivery of Transgenes to Pekin Duck Cells *in vitro* and *in vivo*

**DOI:** 10.1101/2022.03.11.484008

**Authors:** Carleigh R. Robinson, Gregory S. Fraley, Benjamin G. Kopek

## Abstract

The Pekin duck was domesticated between 4000 and 10,000 years ago from the Mallard duck and is the predominate meat type duck in the world (Cherry and Morris, 2008). The global production of waterfowl is a rapidly growing industry. Total meat duck production increased from 2.9 million tons in 2000 to nearly 4.4 million tons in 2013, a growth rate of 3.2% per year, and further increased to 7.2 million tons in 2018, and valued at $19B in 2019 (IndexBox, 2019). Pekin ducks *(Anas platyrhynchus domesticus)* are the fastest growing poultry species growing from hatch to 4.5 kg market weight in as little as 28 days (Blois et al., 2019; Campbell et al., 2015; Cherry and Morris, 2008). Thus, there is a need to study the growth and reproduction of this economically important species. While Pekin duck biology is being explored by many researchers, there are fewer molecular tools available for duck compared to other poultry species and many fewer compared to mammalian systems. For example, one molecular tool commonly used to interrogate other systems are adeno-associated virus (AAV) vectors. AAV vectors are being utilized by many researchers to deliver transgenes to target tissues or cells and for genetic manipulation. Recently, avian adeno-associated virus (A3V) has been used to deliver genes to the cells and neurons of the domestic chicken (*Gallus gallus domesticus*). Here, we show that A3V can be used to deliver genes to Pekin duck neurons and cells in culture. Together, these results suggest that A3V will be a useful molecular tool in Pekin duck research.

**Summary Statement:** Here we demonstrate the use of viral vectors to deliver transgenes to Pekin duck cells. These vectors can be used to advance understanding of reproduction in this economically important species.

## Introduction

While duck meat consumption remains small compared to other poultry species, the global duck meat trade continues to increase and now represents over $19 billion in global trade value (Biswas et al., 2019). Although traditionally associated with Asian countries, duck meat consumption continues to grow in popularity worldwide with the countries of Germany, the United Kingdom, and France among the top five importers of duck meat (Food and Agriculture Organization of the United Nations, 2022). Growth of the duck meat trade is expected to continue to increase by 1.31 billion dollars from 2020-2024 (Technavio).

The rise in duck meat consumption has driven researchers to search for ways to improve reproduction rates to meet the demands of the growing duck meat trade. In many instances, duck continues to be farmed in a traditional manner, but commercial farming of Pekin ducks is a multibillion-dollar industry worldwide (IndexBox, 2019). The top producers of duck meat are China/Myanmar, France, the USA, and the United Kingdom. Among duck species that are farmed for meat, the Pekin duck *(Anas platyrhynchus domesticus)* is the most popular. As such, the Fraley lab (Campbell et al., 2015; Porter et al., 2016; Porter et al., 2018; Potter et al., 2018; Vostrizansky et al., 2022; Wyk and Fraley, 2021) has performed numerous studies on reproduction in the Pekin duck including the characterization of several light sensitive receptors involved in the reproductive axis (Potter et al., 2018). However, relative to the domestic chicken (*Gallus gallus domesticus*) few tools have been tested or developed for probing the neurobiological pathways of Pekin duck.

To develop molecular tools for investigating the neurobiology and reproduction of the Pekin duck, we explored the use of viral vectors. Viral vectors, such as adeno-associated viruses (AAV), lentiviral vectors, and recombinant baculoviruses give scientists the ability to deliver transgenes to a range of organisms (Wang et al., 2019). Adeno-associated viruses (AAV) are in the *Parvoviridae* family of single-stranded DNA viruses. Serotypes of AAV have been found to transduce specific cell types, tissues, and species (Srivastava, 2016). AAV vectors are produced in helper cell lines and the vectors themselves are not competent for replication, thus reducing the chance of pathogenicity. AAV vectors are being tested and used as gene therapy for multiple diseases in humans (He et al., 2021; Wang et al., 2019) including clinical trials of AAV-based gene therapies for cystic fibrosis (Mueller and Flotte, 2008), Alzheimer’s disease (Alves et al., 2016), lipoprotein lipase (LPL) deficiency (Carpentier et al., 2012; Gaudet et al., 2012), and muscular dystrophy (Hinderer et al., 2018; Hordeaux et al., 2018). Luxturna was the first drug utilizing AAV to be approved by the FDA for treatment of Leber congenital amarousis type 2 (Bainbridge et al., 2008; Hauswirth et al., 2008; Maguire et al., 2008). In May 2019, Zolgensma was approved by the FDA to treat spinal muscular atrophy and AAV vectors are being investigated for use in other neurodegenerative diseases like amyotrophic lateral sclerosis (ALS) (Borel et al., 2018; Mendell et al., 2017).

Despite the success of AAV vectors in mammalian systems, AAV vector transduction for many avian species has had limited success. Therefore, we examined the use of avian adeno-associated virus (A3V). A3V was isolated and initially characterized in 1973 (Yates et al., 1973) but was cloned and sequenced in 2003 (Bossis and Chiorini, 2003). A3V is a distinct serotype from primate AAV1-4 (Yates et al., 1973), but there is evidence of A3V infection in humans (Bossis and Chiorini, 2003). Bossis and Chiorini made A3V vectors expressing β-galactosidase and identified the transduction efficiency across multiple cell types (Bossis and Chiorini, 2003). A3V showed little to no transduction in primate cells, but readily transduced multiple primary and immortalized avian cell types (chicken, quail) (Bossis and Chiorini, 2003). Matsui et al. extended this work to show that A3V transduces neurons in chicken (*Gallus gallus domesticus*) brains (Matsui et al., 2012). Furthermore, new transgene constructs expressing fluorescent proteins were made along with inducible promoters (Matsui et al., 2012). In 2019, Waldner et al. demonstrated the ability of A3V to transduce postembryonic chicken retina (Waldner et al., 2019). Thus, A3V is capable of delivering transgenes to both dividing and non-dividing avian cell lines.

To extend the work of others and advance our own interests in Pekin duck research, we examined the ability of A3V to transduce Pekin duck cells in culture and *in vivo*. We found that A3V can effectively transduce Pekin duck cells in culture and express fluorescent transgenes. Additionally, A3V was able to transduce neurons in adult Pekin duck brains. Thus, A3V can transduce both dividing and non-dividing Pekin duck cells. This work demonstrates that A3V may be a valuable tool to further goals related to research in Pekin duck reproduction.

## Methods

### Cell culture

HEK 293T cells were a gift from Kristin Dittenhaffer-Reed (Hope College, USA) and grown in DMEM (ThermoFisher Sci. #11965092) with 10% FBS (ThermoFisher Sci. #A3160501) at 37 °C and 5% CO_2_. Pekin duck (*Anas platyrhynchus domesticus*) cells were obtained from ATCC (CCL-141) and grown in DMEM (ThermoFisher Sci. #11965092) with 10% FBS (ThermoFisher Sci. #A3160501) at 37 °C and 5% CO_2_.

### Plasmids and molecular cloning

pA3V-RSV-mCherry, pA3V-RSV-GFP, pA3V Rep/Cap plasmids were gifts from Dr. Dai Watanabe (Kyoto University, Japan) with permission from Dr. Chorini (NIH, USA). The helper plasmid pAd-Helper (pAd-DeltaF6) was a gift from James M. Wilson (Addgene plasmid #112867; http://n2t.net/addgene:112867 ; RRID:Addgene_112867).

### Virus Production

A3V vectors were produced using common methods and described previously (Matsui et al., 2012). Briefly, A3V vectors were produced through transfection of HEK 293T cells with the three plasmids pAd-helper, pA3VRep/Cap, and the pA3V-RSV-mCherry or pA3V-RSV-eGFP. *Trans*IT-VirusGEN (MirusBio #MIR 6703) was used as the transfection reagent. Three days post-transfection, A3V was harvested and purified from the HEK293FT cells. To ensure complete lysis of the HEK293FT cells, the cell and media mixture was frozen completely in a dry ice/ethanol bath followed by a 37 °C water bath to thaw the cells. This process was repeated a total of three times for each sample. Next, benzonase (MilliporeSigma #E8263) was added to the cell media mixture in a 50 U/mL concentration. Following centrifugation, the cell debris was discarded appropriately and the virus-containing lysate was kept for further purification.

Purification of genome containing virions (as opposed to empty capsids) was done using an isomolar density gradient medium (iodixanol). The gradient consisted of four density steps (15%, 25%, 40%, 60%). The cell lysate was added to the top of the gradient and centrifuged at 70,000 rpm for 75 min at 10 °C in a T70i rotor. Following ultracentrifugation, fractions (0.5 mL to 1.0 mL) were collected from the gradient. The genome containing virions were expected to be at the interface between the 40% and 60% solutions. The collected fractions were examined by protein gel to identify fractions containing proteins of the expected molecular weights of the three A3V capsid proteins. These fractions were pooled and concentrated using centrifugal filters (EMDMillipore Amicon Ultra-15 #UFC910008; cellulose 100,000 NMWL). This also included a buffer exchange with PBS to remove iodixanol.

### Genomic Copy Determination of A3V Vectors

There is no suitable plaque assay for adeno-associated viruses so quantitative PCR (qPCR) was used to determine the genome copies per mL (gc/mL). A plasmid was used as a standard for calculation.

### A3V Transduction of Pekin Duck Cells in Culture

Pekin duck (*Anas platyrhynchus domesticus)* embryo cells (ATCC, # CCL-141) were plated into the wells of a chambered coverglass (MilliporeSigma #Z734853) at a density of 10,000 cells/well. Cells were infected at an MOI of 100 based on the gc/ML determined from qPCR. The cells were imaged using a Nikon A1^+^ confocal fluorescence microscope 24-72 hours post infection.

### A3V Transduction of Pekin Duck Neuronal Cells *in vivo*

The A3V-RSV-EGFP vector was used for stereotaxic microinjections into the lateral ventricle of the Pekin Duck brain. Adult (n = 6) adult female ducks (∼4.0 kg) were anesthetized with proprofol anesthesia injected intravenously (10mg/ml approximately 5 ml induction and additional to effect). Prior to surgery, ducks were injected intramuscularly with carprofen as an analgesic (2mg/ml/kg). Ducks were placed in a Kopf stereotaxic atlas with their heads declined at 45° angle in order to place the brain in a horizontal position. The skin was incised and fascia removed from skull. A trephine hole was drilled (AP = -10mm to periorbita, +1mm off midline) and a Hamilton microliter syringe (30 Ga needle) was lowered to the depth of the lateral ventricle (−3.0mm). Injection was made over a 5-minute time period, and needle held in place for 2 minutes following injection. The needle was removed and trephine hole filled with bone wax, and skin closed with suture glue. Each injection utilized 5 µl of the designated vector at a titer of 7.06 × 10^10^ genomic copies/mL. The animals recovered following injections and neuronal cell samples were taken and imaged ten days after injections.

To image brain sections, ducks were euthanized using pentobarbital (IV, 396 mg/ml/kg). Brains were rapidly removed and immediately frozen on dry ice, and stored at -80°C until processed. Frozen sections of duck brain were collected at 40 μm in the coronal plane and direct mounted on Superfrost Plus™ slides, and allowed to air dry. Sections were collected form the septomesencephalic tract rostrally to the third cranial nerve caudally. This region spans the diencephalon with the injection site at its center. Slides were coverslipped using Vectashield Hardset media without DAPI. Slide analyses were performed on a Leica DM350 fluorescent microscope and a Nikon A1^+^ confocal fluorescence microscope.

## Results

Many serotypes of adeno-associated virus capsids exist that permit the targeting of the vectors to specific cell types. To target avian cell types specifically, the avian-adeno associated virus capsid was delivered by the pA3V-Rep/Cap plasmid (Figure 1A). (Matsui et al., 2012). The transgene expression plasmid consists of A3V inverted terminal repeats (ITR) that flank an expression cassette that is packaged into the capsid (Figure 1A). A Rous sarcoma virus (RSV) promoter is used because it is known to drive high expression levels of transgenes in many species (Gorman et al., 1982). The common pAd-Helper plasmid was used for vector production in HEK 293T cells. Vector production consisted of a triple transfection of the expression and packaging plasmids into HEK293T cells (Figure 1B). A3V was harvested and empty capsids removed using density gradient ultracentrifugation (see Methods). Quantitative PCR indicated the A3V-RSV-EGFP vector preparation had 7.06 × 10^10^ gc/mL and the A3V-RSV-mCherry preparation contained 3.30 × 10^10^ gc/mL.

**Figure 1.**
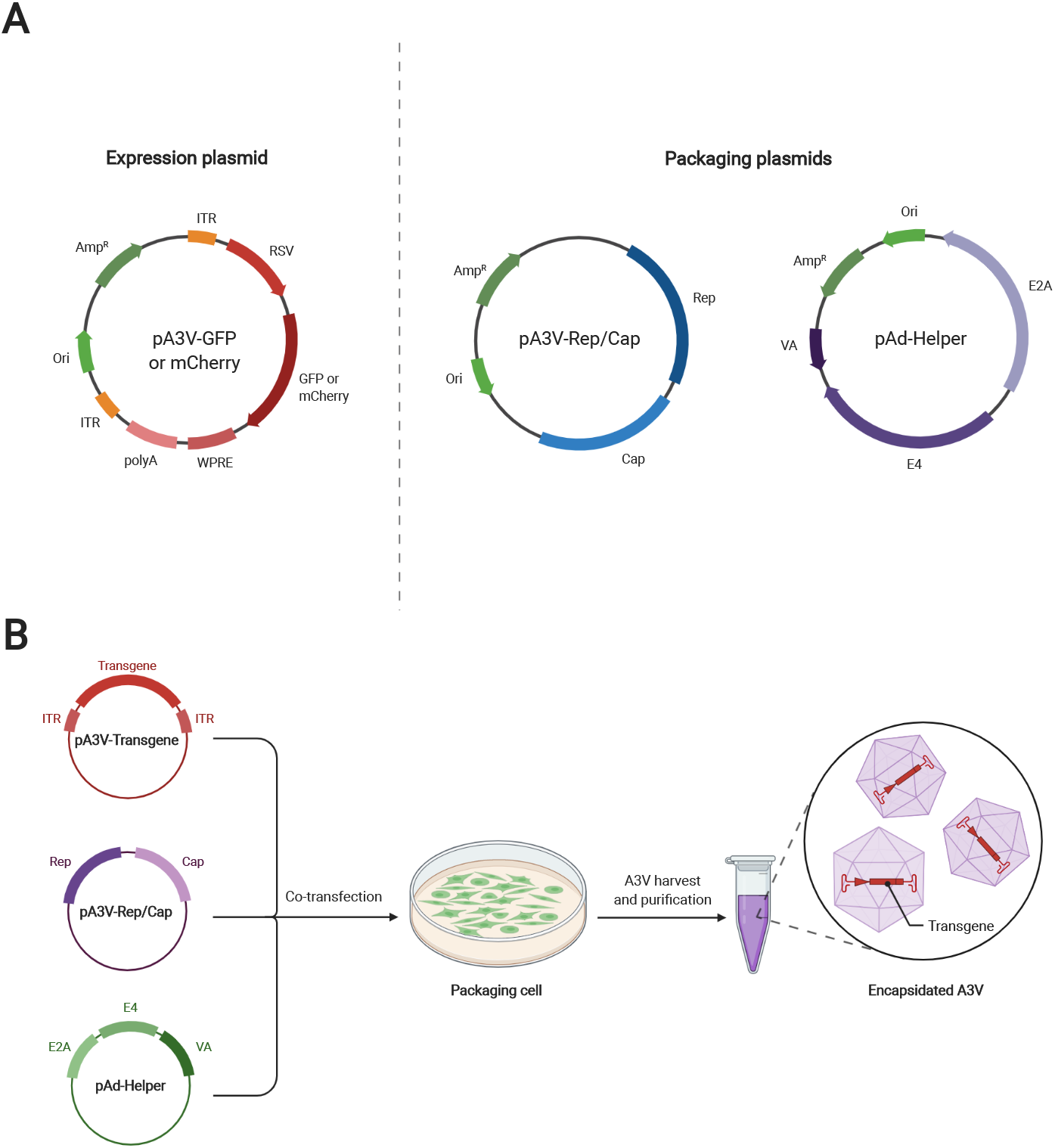
Avian adeno-associated vector production. (A) General features of plasmids used for A3V vector production. In the expression plasmid, a Rous sarcoma virus (RSV) promoter drives expression of a fluorescent protein (GFP or mCherry). The inverted terminal repeats (ITR) are avian adeno-associated virus specific. WPRE; woodchuck hepatitis virus posttranscriptional regulatory element, polyA; polyadenylation signal sequence from simian virus 40, Ori; bacterial origin of replication, Amp^R^; ampicillin resistance gene. (Right of dashed line) Packaging plasmids for A3V vector production. pA3V-Rep/Cap expresses the A3V replication and capsid proteins. pAd-Helper encodes adenoviral proteins involved in virus production. (B) General scheme for A3V vector production. All three plasmids from (A) are transfected into the packaging cell line HEK-293T. A3V is then purified from these cells and genome copies (indicating infectious virions) are determined using qPCR. (A) Adapted from “Plasmids for AAV Vector Production” and (B) adapted from “AAV Production by Triple Transfection”, BioRender.com (2022). Retrieved from https://app.biorender.com/biorender-templates.

To determine if A3V is able to transduce Pekin duck cell lines, we infected a Pekin duck embryonic cell line with the fluorescent transgene expressing A3V vectors (Figure 2). Some fluorescence was observed at 24 hours post infection and continued to increase up to 72 hours post infection. Fluorescent protein expression plateaued at 72 hours post infection with stable expression continuing to seven days in culture. Transduced cells appeared to remain healthy during the course of the experiment (Figure 2, center column). These data indicate A3V is able to transduce Pekin duck cells *in vitro* with little impact on cell viability.

**Figure 2.**
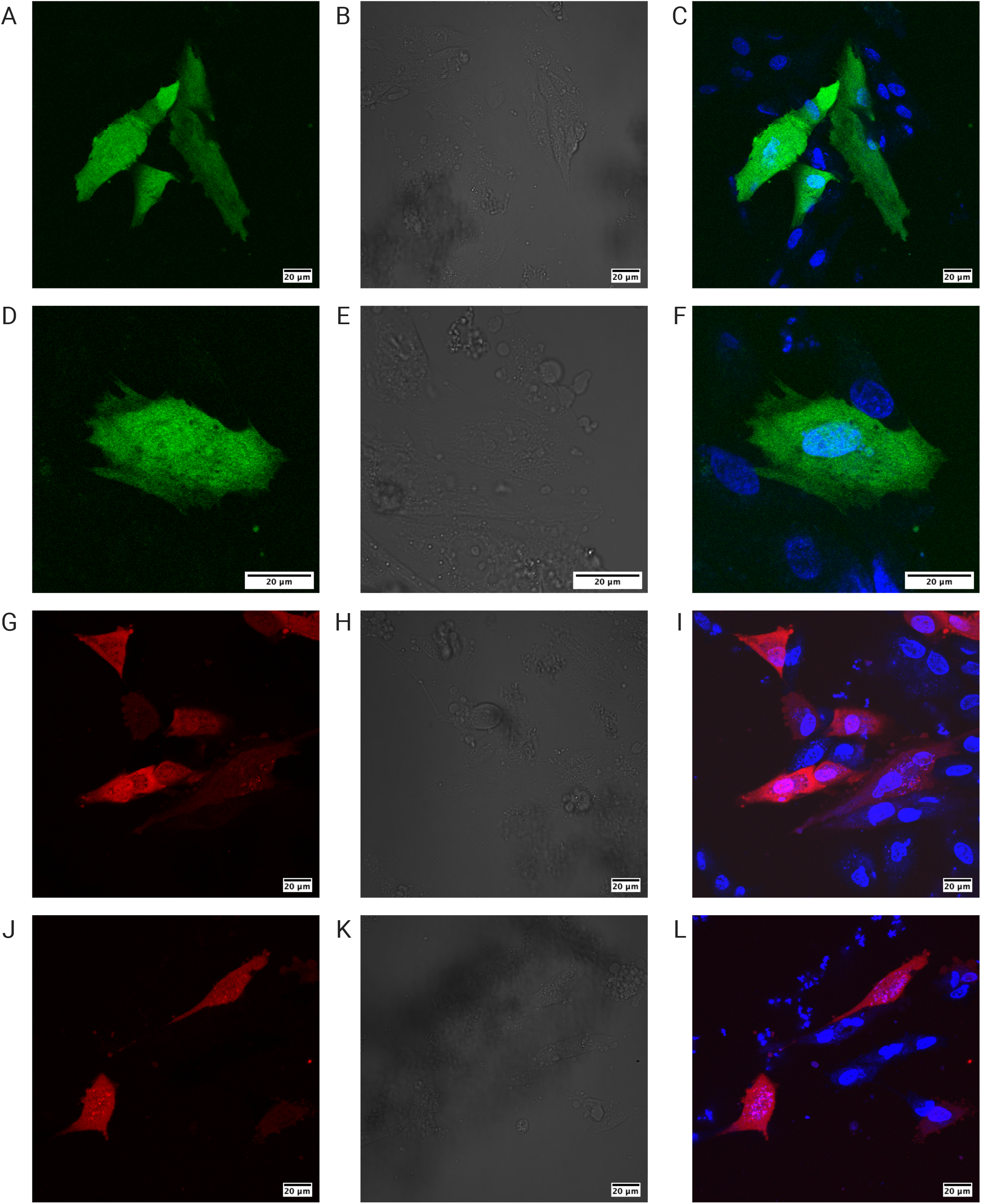
Infection of Pekin duck cells in culture demonstrates infection with A3V vectors and transgene expression. Pekin duck cells grown in culture were infected with A3V vectors expressing either GFP (A-F) or mCherry (G-L) transgenes. Infected cells were imaged using confocal fluorescence microscopy. Left column shows expression of the transgene (GFP; A, D or mCherry; G, J). Middle column shows the brightfield images of the same area. Right column shows a merged signal of the fluorescent transgene signal with a DNA stain (Hoescht) indicating the nucleus.

To test whether the vectors can deliver transgenes *in vivo*, Pekin duck were injected with A3V containing the GFP expression cassette (Figure 3). Ten days post injection, the animals were sacrificed and the brain tissue was harvested and sectioned. Confocal fluorescence microscopy was used to examine the tissue slices for GFP expression. When areas near the injection site (* in Figure 3) were examined, several cells with neuronal morphology were found to be expressing GFP. Using the z-sectioning capabilities of the confocal microscopy, long neuronal axons can be visualized for several of the cells expressing GFP. Thus, A3V vectors can transduce Pekin duck cells with an apparent preference for neurons. This preference for neurons is consistent with data from chicken (Matsui et al., 2012).

**Figure 3.**
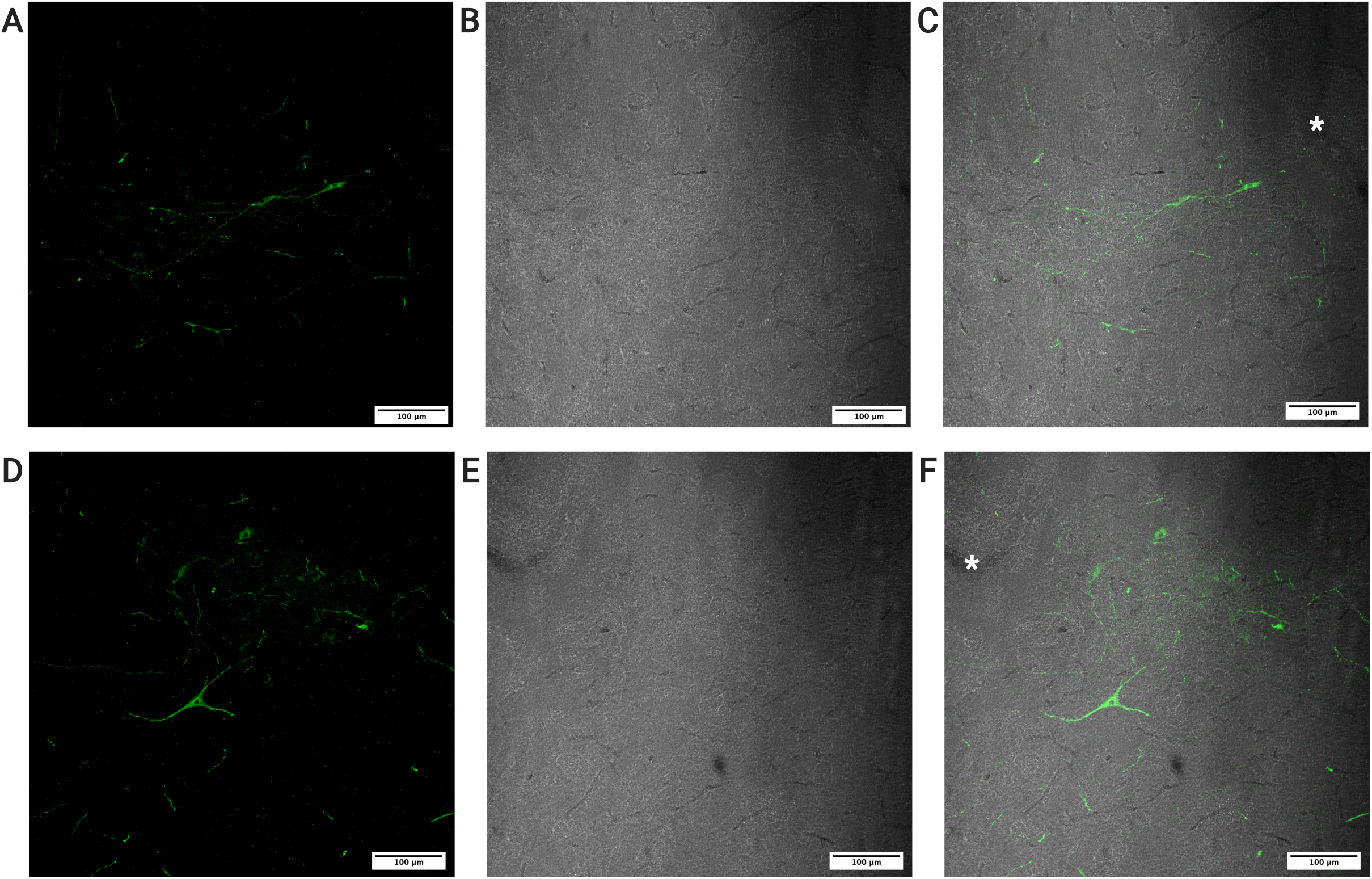
Infection of Pekin duck brain tissue with A3V vectors demonstrate transgene expression in neural tissue. Pekin duck were injected with a solution containing the A3V vector carrying the GFP transgene (see Methods). Brain tissue was harvested and a z-stack of images were collected by confocal fluorescence microscopy. A maximum intensity projection was created from the z-stack. Panels A and D show GFP expression in neural cells. Panels B and E show the brightfield image of the tissue and panels C and F are a merge of the fluorescent and brightfield channels. *indicates injection site.

## Discussion

The purpose of this study was to develop a molecular tool that can be used to transiently alter gene expression in duck brains. Such a tool would greatly improve the neuroendocrine research on growth and reproduction in this economically important species. We demonstrated that A3V vectors can be used to deliver exogenous genes to Pekin duck cells *in vitro* and *in vivo*. To our knowledge, this is the first example of viral mediated transgene delivery to Pekin duck cells in culture and *in vivo*. Although using adeno-associated viral vectors has been done for other avian species, duck has not been tested (Bossis and Chiorini, 2003; Matsui et al., 2012). Recombinant baculoviruses have been used to transduce avian cells lines but in a non-cell-type specific manner and only in primary cultures (Song et al., 2006). Therefore, these A3V vectors will be important tools to move research forward in this economically important species.

The use of AAV vectors has been vital for many neuroscience studies. For other organisms, recombinant AAV vectors have been used for neural circuit analysis (Betley and Sternson, 2011). Targeted injection of A3V vectors expressing these fluorescent transgenes could be used to dissect neural circuits in Pekin duck. A3V vectors could also be used to manipulate neural circuitry by targeting transgenes that activate or silence defined neural populations. Chemogenetic tools, such as DREADDs (Designer Receptors Exclusively Activated by Designer Drugs) can provide excitatory or inhibitory control over neuronal function (Roth, 2016).

Another opportunity is gene knockout using CRISPR/Cas9 systems. Although A3V vectors have limiting packaging capacity, there have been ways developed to package the elements required for CRISPR into AAV (Li et al., 2018; VanDusen et al., 2017), allowing for specific gene editing in neurons (Kumar et al., 2018). The Pekin duck genome has recently been sequenced (Huang et al., 2013; Li et al., 2021), which is essential for targeted and specific gene knockouts. Our lab has already generated the A3V expression cassettes containing the SaCas9 endonuclease and will soon begin designing sgRNA sequences to target the genetic elements involved in Pekin duck reproduction. In future studies, we will use this system in an attempt to better understand the roles of the various factors in Pekin duck reproduction.

## Acknowledgements

We thank Dr. S. Fraley for help with Pekin duck anesthesia. HEK 293T cells were a gift from Dr. K. Dittenhafer-Reed (Hope College, Holland, MI). We thank Dr. Dai Watanabe (Kyoto University, Japan) for plasmids pA3V-RSV-mCherry, pA3V-RSV-GFP, pA3V Rep/Cap. The helper plasmid pAd-Helper (pAd-DeltaF6) was a gift from James M. Wilson (Addgene plasmid #112867; http://n2t.net/addgene:112867 ; RRID:Addgene_112867).

## References

Alves, S., Fol, R. and Cartier, N. (2016). Gene Therapy Strategies for Alzheimer’s Disease: An Overview. Hum Gene Ther 27, 100–107.

Bainbridge, J. W. B., Smith, A. J., Barker, S. S., Robbie, S., Henderson, R., Balaggan, K., Viswanathan, A., Holder, G. E., Stockman, A., Tyler, N., et al. (2008). Effect of gene therapy on visual function in Leber’s congenital amaurosis. New Engl J Medicine 358, 2231–9.

Betley, J. N. and Sternson, S. M. (2011). Adeno-Associated Viral Vectors for Mapping, Monitoring, and Manipulating Neural Circuits. Hum Gene Ther 22, 669–677.

Biswas, S., Banerjee, R., Bhattacharyya, D., Patra, G., Das, A. K. and Das, S. K. (2019). Technological investigation into duck meat and its products - a potential alternative to chicken. World’s Poult Sci J 75, 609–620.

Blois, L. V., Bentley, A., Porter, L., Prihoda, N., Potter, H., Wyk, B. V., Shafer, D., Fraley, S. M. and Fraley, G. S. (2019). Feed Restriction Can Alter Gait but Does not Reduce Welfare in Meat Ducks. J Appl Poultry Res 28, 858–866.

Borel, F., Gernoux, G., Sun, H., Stock, R., Blackwood, M., Brown, R. H. and Mueller, C. (2018). Safe and effective superoxide dismutase 1 silencing using artificial microRNA in macaques. Sci Transl Med 10, eaau6414.

Bossis, I. and Chiorini, J. A. (2003). Cloning of an Avian Adeno-Associated Virus (AAAV) and Generation of Recombinant AAAV Particles. J Virol 77, 6799–6810.

Campbell, C. L., Colton, S., Haas, R., Rice, M., Porter, A., Schenk, A., Meelker, A., Fraley, S. M. and Fraley, G. S. (2015). Effects of different wavelengths of light on the biology, behavior, and production of grow-out Pekin ducks. Poultry Sci 94, 1751–1757.

Carpentier, A. C., Frisch, F., Labbé, S. M., Gagnon, R., Wal, J. de, Greentree, S., Petry, H., Twisk, J., Brisson, D. and Gaudet, D. (2012). Effect of Alipogene Tiparvovec (AAV1-LPL S447X) on Postprandial Chylomicron Metabolism in Lipoprotein Lipase-Deficient Patients. J Clin Endocrinol Metabolism 97, 1635–1644.

Cherry, P. and Morris, T. (2008). Domestic duck production: science and practice. (ed. Cherry, P.) and Morris, T.) CABI. Food and Agriculture Organization of the United Nations (2022).

Gaudet, D., Méthot, J., Déry, S., Brisson, D., Essiembre, C., Tremblay, G., Tremblay, K., Wal, J. de, Twisk, J., Bulk, N. van den, et al. (2012). Efficacy and long-term safety of alipogene tiparvovec (AAV1-LPLS447X) gene therapy for lipoprotein lipase deficiency: an open-label trial. Gene Ther 20, 361–369.

Gorman, C. M., Merlino, G. T., Willingham, M. C., Pastan, I. and Howard, B. H. (1982). The Rous sarcoma virus long terminal repeat is a strong promoter when introduced into a variety of eukaryotic cells by DNA-mediated transfection. Proc National Acad Sci 79, 6777–6781.

Hauswirth, W. W., Aleman, T. S., Kaushal, S., Cideciyan, A. V., Schwartz, S. B., Wang, L., Conlon, T. J., Boye, S. L., Flotte, T. R., Byrne, B. J., et al. (2008). Treatment of leber congenital amaurosis due to RPE65 mutations by ocular subretinal injection of adeno-associated virus gene vector: short-term results of a phase I trial. Hum Gene Ther 19, 979–90.

He, X., Urip, B. A., Zhang, Z., Ngan, C. C. and Feng, B. (2021). Evolving AAV-delivered therapeutics towards ultimate cures. J Mol Medicine Berlin Ger 99, 593–617.

Hinderer, C., Katz, N., Buza, E. L., Dyer, C., Goode, T., Bell, P., Richman, L. K. and Wilson, J. M. (2018). Severe Toxicity in Nonhuman Primates and Piglets Following High-Dose Intravenous Administration of an Adeno-Associated Virus Vector Expressing Human SMN. Hum Gene Ther 29, 285–298.

Hordeaux, J., Wang, Q., Katz, N., Buza, E. L., Bell, P. and Wilson, J. M. (2018). The Neurotropic Properties of AAV-PHP.B Are Limited to C57BL/6J Mice. Mol Ther 26, 664–668.

Huang, Y., Li, Y., Burt, D. W., Chen, H., Zhang, Y., Qian, W., Kim, H., Gan, S., Zhao, Y., Li, J., et al. (2013). The duck genome and transcriptome provide insight into an avian influenza virus reservoir species. Nat Genet 45, 776–783.

IndexBox (2019). Global Duck And Goose Meat Market to Keep Growing, Driven by Strong Demand in Asia - Global Trade Magazine.

Kumar, N., Stanford, W., Solis, C. de, Aradhana Abraham, N. D., Dao, T.-M. J., Thaseen, S., Sairavi, A., Gonzalez, C. U. and Ploski, J. E. (2018). The Development of an AAV-Based CRISPR SaCas9 Genome Editing System That Can Be Delivered to Neurons in vivo and Regulated via Doxycycline and Cre-Recombinase. Front Mol Neurosci 11, 413.

Li, A., Lee, C. M., Hurley, A. E., Jarrett, K. E., Giorgi, M. D., Lu, W., Balderrama, K. S., Doerfler, A. M., Deshmukh, H., Ray, A., et al. (2018). A Self-Deleting AAV-CRISPR System for In Vivo Genome Editing. Mol Ther Methods Clin Dev 12, 111–122.

Li, J., Zhang, J., Liu, J., Zhou, Y., Cai, C., Xu, L., Dai, X., Feng, S., Guo, C., Rao, J., et al. (2021). A new duck genome reveals conserved and convergently evolved chromosome architectures of birds and mammals. Gigascience 10, giaa142.

Maguire, A. M., Simonelli, F., Pierce, E. A., Pugh, E. N., Mingozzi, F., Bennicelli, J., Banfi, S., Marshall, K. A., Testa, F., Surace, E. M., et al. (2008). Safety and Efficacy of Gene Transfer for Leber’s Congenital Amaurosis. New Engl J Med 358, 2240–2248.

Matsui, R., Tanabe, Y. and Watanabe, D. (2012). Avian adeno-associated virus vector efficiently transduces neurons in the embryonic and post-embryonic chicken brain. PloS One 7, e48730.

Mendell, J. R., Al-Zaidy, S., Shell, R., Arnold, W. D., Rodino-Klapac, L. R., Prior, T. W., Lowes, L., Alfano, L., Berry, K., Church, K., et al. (2017). Single-Dose Gene-Replacement Therapy for Spinal Muscular Atrophy. New Engl J Medicine 377, 1713–1722.

Mueller, C. and Flotte, T. R. (2008). Clinical gene therapy using recombinant adeno-associated virus vectors. Gene Therapy 15, 858–863.

Porter Alenciks, E. and Fraley, G. (2016). Gonadal regression elicited in Pekin duck drakes and hens associated with supplemental light from kerosene lanterns during the winter months. In Poultry Science Association Annual Meeting, p. Abstract #195.

Porter, L., Porter, A., Potter, H., Alenciks, E., Fraley, S. M. and Fraley, G. S. (2018). Low light intensity in Pekin duck breeder barns has a greater impact on the fertility of drakes than hens. Poultry Sci 97, 4262–4271.

Potter, H., Alenciks, E., Frazier, K., Porter, A. and Fraley, G. S. (2018). Immunolesion of melanopsin neurons causes gonadal regression in Pekin drakes (Anas platyrhynchos domesticus). General and Comparative Endocrinology 256, 16–22.

Roth, B. L. (2016). DREADDs for Neuroscientists. Neuron 89, 683–694.

Song, J., Liang, C. and Chen, X. (2006). Transduction of avian cells with recombinant baculovirus. J Virol Methods 135, 157–162.

Srivastava, A. (2016). In vivo tissue-tropism of adeno-associated viral vectors. Curr Opin Virol 21, 75–80.

Technavio Global Duck Meat Market to Grow by $ 1.31 Billion During 2020-2024 | Featuring AJC International Inc., AMI LLC sp.k, and BRF SA Among Others | Technavio | Business Wire.

VanDusen, N. J., Guo, Y., Gu, W. and Pu, W. T. (2017). CASAAV: A CRISPR-Based Platform for Rapid Dissection of Gene Function In Vivo. Curr Protoc Mol Biology 120, 31.11.1-31.11.14.

Vostrizansky, A., Barce, A., Gum, Z., Shafer, D. J., Jeffrey, D., Fraley, G. S. and Rivera, P. D. (2022). Effect of pre-hatch incubator lights on the ontogeny of CNS opsins and photoreceptors in the Pekin duck. Poultry Sci 101, 101699.

Waldner, D. M., Visser, F., Fischer, A. J., Bech-Hansen, N. T. and Stell, W. K. (2019). Avian Adeno-Associated Viral Transduction of the Postembryonic Chicken Retina. Transl Vis Sci Technology 8, 1.

Wang, D., Tai, P. W. L. and Gao, G. (2019). Adeno-associated virus vector as a platform for gene therapy delivery. Nat Rev Drug Discov 18, 358–378.

Wyk, B. V. and Fraley, G. (2021). Ontogeny of OPN4, OPN5, GnRH and GnIH mRNA Expression in the Posthatch Male and Female Pekin Duck (Anas platyrhynchos domesticus) Suggests OPN4 May Have Additional Functions beyond Reproduction. Animals 11, 1121.

Yates, V. J., Mishad, A. M. E., McCormick, K. J. and Trentin, J. J. (1973). Isolation and Characterization of an Avian Adenovirus-Associated Virus. Infect Immun 7, 973–980.

